# A scalable and robust system for Audience EEG recordings

**DOI:** 10.1101/2022.12.16.520764

**Authors:** Georgios Michalareas, Ismat M.A. Rudwan, Claudia Lehr, Paolo Gessini, Alessandro Tavano, Matthias Grabenhorst

**Author notes:** CORRESPONDENCE TO: Georgios Michalareas, Max Planck Institute for Empirical Aesthetics, Grueneburgweg 14, 60322 Frankfurt am Main, Germany. Contact.

## Abstract

The neural mechanisms that unfold when humans form a large group defined by an overarching context, such as audiences in theater or sports, are largely unknown and unexplored. This is mainly due to the lack of availability of a scalable system that can record the brain activity from a significantly large portion of such an audience simultaneously. Although the technology for such a system has been readily available for a long time, the high cost as well as the large overhead in human resources and logistic planning have prohibited the development of such a system. However, during the recent years reduction in technology costs and size have led to the emergence of low-cost, consumer-oriented EEG systems, developed primarily for recreational use. Here by combining such a low-cost EEG system with other off-the-shelve hardware and tailor-made software, we develop in the lab and test in a cinema such a scalable EEG hyper-scanning system. The system has a robust and stable performance and achieves accurate unambiguous alignment of the recorded data of the different EEG headsets. These characteristics combined with small preparation time and low-cost make it an ideal candidate for recording large portions of audiences.

**Highlights:** A scalable EEG hyper-scanning system for recording audiences and large groups is presented with the following characteristics.

- Off-the-shelve, low cost components, namely a MUSE EEG headset, a Raspberry Pi computer and Photodiode.
- A Python library, available to the public, has been specifically developed from first principles, optimized for facilitating robust recording over Bluetooth even when multiple EEG headsets are in close proximity.
- The use of photodiodes provides unambiguous data alignment between the different systems.
- A proof-of-concept system with 10 EEG headsets has been tested in the lab but also in naturalistic conditions, recording members of the audience in four different long movie screenings in the cinema of the German Film Museum.

## INTRODUCTION

“We human beings are social beings. We come into the world as the result of others’ actions. We survive here in dependence on others. Whether we like it or not, there is hardly a moment of our lives when we do not benefit from others’ activities. For this reason, it is hardly surprising that most of our happiness arises in the context of our relationships with others.” Dalai Lama XIV

This quote by Dalai Lama highlights characteristically the role of one of the most defining factors of human behavior, the ever-present interaction with other humans to a lesser or higher degree and at a direct or indirect level. This very important factor, which dominates our behavior and our brain function in everyday life has largely been ignored in neuroscience research until recently. Most of the Neuroscience research has been focused on the single brain, studied in isolation within the controlled conditions of a laboratory.

During the last two decades there has been a sharp increase in the number of studies in which brain activity of two or more interacting humans is recorded simultaneously. This type of experiments are usually termed “hyper-scanning”. The main neuroscience modalities used in such experiments are Electroencephalography (EEG), Magnetoencephalography (MEG), functional Near-Infrared Spectroscopy(fNIRS) and functional MRI (fMRI). fMRI does not measure directly neural activity but measures changes in blood flow associated with it. It has the highest spatial accuracy but low temporal resolution. Due to the fact that only one person fits inside an MRI scanner, hyper-scanning experiments can only occur through indirect interaction. It is the least naturalistic hyper-scanning modality. fNIRS similarly measures brain activity indirectly through light refraction changes due to changes in hemoglobin in the cortical surface. It has a low spatial and low temporal resolution. Its advantage is that it highly mobile and is relatively immune to movement artifacts. MEG has a very high temporal resolution and modest spatial resolution. Its main disadvantage is that due to the structural characteristics of the MEG scanner, hyper-scanning cannot be performed directly but indirectly, similar to fMRI. Finally, EEG has a very high temporal resolution but modest to low spatial resolution. Its main advantage is that it can be highly mobile and that with even few electrodes brain activity from a large part of the human brain can be recorded. EEG and fNIRS have been the most popular hyper-scanning modalities during the last decade. Due to its high temporal resolution, EEG is the main method of hyper-scanning when the fine temporal dynamics of interbrain interactions are the focus of research. This is also the main focus of this study.

The vast majority of EEG hyper-scanning experiments have studied interactions in pairs. A long list of such experiments has covered various aspects of interactive brain dynamics such as joint action, shared attention, interactive decision making, affective communication and others. Recent reviews offer a very extensive insight of these studies (Barde et al., 2020; Czeszumski et al., 2020; Liu et al., 2018; Schilbach et al., 2013). A few EEG studies went further than pairs and studied interbrain interactions between three or four musicians using laboratory-grade EEG headsets (Babiloni et al., 2012; Babiloni et al., 2011; Müller et al., 2018).

One of the main limiting factors of scaling up the number of simultaneously scanned participants has been the typically high cost of laboratory-grade EEG systems. The purchase of multiple such systems entails high cost while the research questions tackled are mostly exploratory and only very few institutions have the capacity to fund such high-cost, high-risk type of research. However, in the last decade various consumer-grade EEG systems, with low cost, appeared in the market mainly for recreational use. Such systems have a small number of electrodes and lower quality of recorded signal. The research community recognized the potential of such cost-effective systems and various studies investigated their suitability for research. (Badcock et al., 2013; Krigolson et al., 2017; LaRocco et al., 2020; Ratti et al., 2017). In all these studies the main outcome was that these systems, despite their lower quality, still provide meaningful recordings that can be used to address research questions.

Two of the most cost-effective and thus popular such systems are the EMOTIV EPOC (EMOTIV, 2022) and MUSE (InteraXon, 2022). The EPOC headset has 14 saline-based electrodes, placed on flexible arms covering a large area of the head. This headset requires significant preparation effort to place and wet the saline sponges on the electrode contacts and subsequently make sure that the electrodes touch the surface on the head of the participant. With elapsed time the quality of the signal drops as the saline solution dries out. Slight movements can cause the flexible arms to move and this causes the electrodes to lose contact. Despite these issues, this headset has been used in many studies. Regarding hyper-scanning it was first used to record a very large number of pairs in an art installation project (Dikker et al., 2021). The two most innovative studies with this headset involved the recording of 12 people simultaneously and the study of inter-brain synchrony. In one study these 12 people were school students attending different types of teaching (Dikker et al., 2017). In the other study 12 members of the audience were recorded during a music festival and their interbrain synchrony was correlated with their experience as audience (Chabin et al., 2022). These studies were proofs-of-concept that hyper-scanning of larger numbers of people is feasible without a prohibitive cost. Although these studies served well as proofs-of-concept, they did not provide a system design that can be easily scalable to much larger numbers, so that potentially an entire audience can be scanned. This is primarily due to the significant time required for preparation (due to the sponges, the saline solution, the corresponding necessary cleaning and the need for adjusting each electrode arm so that the electrode sponges touch the skin through the hair) but also due to the proprietary per-use pricing model. Both of these induce significant barrier in scalability to large numbers.

The MUSE EEG headset has 4 EEG sensors, 2 anterior frontal and 2 temporo-parietal (Fig 1A). The electrodes are dry and the headset is designed so that it can be very quickly placed on the head of the participants and then they can further adjust it by themselves. So the preparation time is very little. The electrodes have been placed in locations where contact is preserved on the head even in long recordings. So in terms in participant preparation this headset is suitable for scaling up the number of recorded people. In terms of recored brain signal quality, MUSE has been shown to be able to capture Event Related Potentials (ERPs) (Krigolson et al., 2021; Krigolson et al., 2017). It has also been shown to be able various aspects of induced and intrinsic brain activity in various aspects of human behavior, such Stress (Asif et al., 2019; Phutela et al., 2022), Attention(Vortmann et al., 2022), Meditation (Kim et al., 2022; Sharma et al., 2022), Mindfulness (Hawley et al., 2021; Hunkin et al., 2021), Rapid diagnosis of stroke (Wilkinson et al., 2020), Drowsiness (LaRocco et al., 2020), Emotion Classification (Raheel et al., 2019) and Fatigue(Krigolson et al., 2021; Ruyi et al., 2017). In a recent study MUSE headsets were used in a hyper-scanning study in an art installation (Chen et al., 2022), which was previously using EMOTIV EPOC headsets (Dikker et al., 2021). Results seemed to be comparable with both headsets.

**Figure 1.**
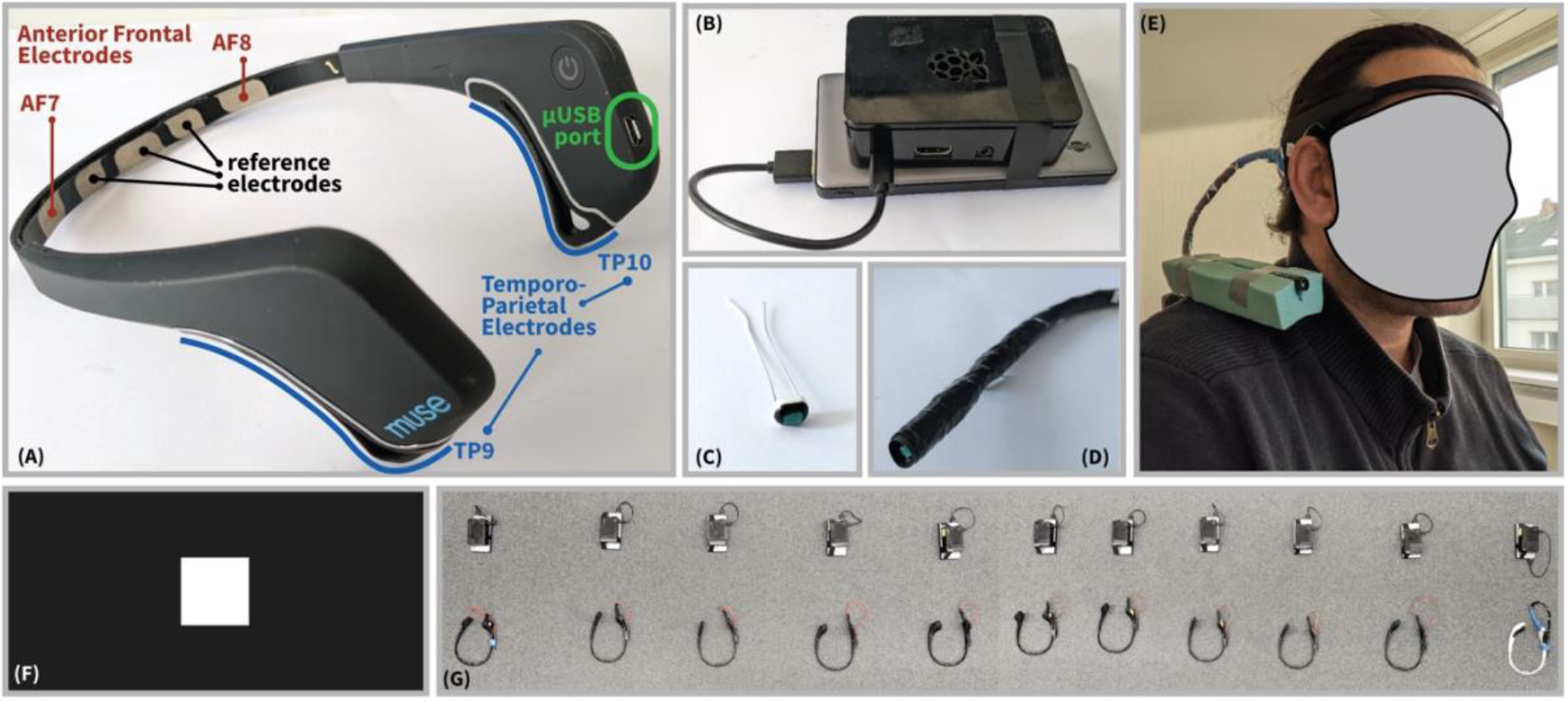
Testing MUSE in the Lab. A) Electrodes of the MUSE headset. There are 2 Anterior Frontal electrodes, AF7 and AF8, on the forehead and two Temporoparietal electrodes, TP9 and TP10, located behind the ears. Notice that TP9 and TP10 are long conductive rubber electrodes, running along the back of the ear. There is one auxiliary electrode channel, accessible through pin 4 of the micro-USB port, located at the right posterior side of the MUSE and in close proximity to electrode TP10. (B) The MUSE headset was recorded through Bluetooth in a Raspberry Pi 3+ which was powered by a lithium battery. (C) A photodiode was tested as a potential sensor for providing signals suitable for aligning different MUSE headsets. The specific photodiode was a planar photodiode, filter-coated for approximating the response of the human eye (see Methods). (D) The photodiode was connected to a micro-USB connector through twisted-wires for minimizing contamination of the other electrodes by the electromagnetic field produced by the photodiode’s current. The photodiode was further insulated with aluminum conductive tape. (E) In the on-head testing the photodiode was attached to a piece of polyethylene terephthalate foam and taped on the shoulder of the participant in order to avoid taping it on the headset and thus potentially introducing electromagnetic contamination to the other electrodes. (F) The testing stimulus was a white square in the center of screen on a black background. The square alternated between black and white with various durations (See Methods). (G) The 10 different MUSE-RPi sets were placed on the floor along a line, in close proximity (~40cm), resembling the distances of people seated in a row in an audience.

Given that MUSE can measure various aspects of brain function, that it requires very small participant preparation time, that its electrodes remain stable in position even in long intervals and is very cost effective, it is a suitable headset for a scalable hyper-scanning system that can record large numbers of participants. Such a system would enable recording large groups such as crowds and audiences without a prohibitive cost of necessary human and technical resources. Additionally, such a system would also allow laboratories and researchers with limited budget resources to do research with hyper-scanning on interbrain dynamics with a relatively small cost. Finally, such a system would be ideal for school laboratories, to assist students in learning about the functional principles of the brain and instigate in them curiosity and motivation towards scientific research.

The case of audiences is particularly of interest, especially in sports and theater where the role of the audience is integral to and vital for the performance and is irreducible to the sum of the experience of each individual in isolation. Very little is known about the neural processes that are partaking in such behavior and one of the main reasons is that up to know a scalable system that can record the brain activity in a large audience has not been available due to necessary high cost and human resources overhead. However, in the last decade the necessary technology for building such a system has become increasingly more available.

In the current study we present the development and testing of a proof-of-concept of such scalable hyper-scanning system. It is based on off-the-shelve components, namely a MUSE EEG headset, a Raspberry Pi computer (RPi) and a photodiode. Each RPi is dedicated to recording over Bluetooth a specific MUSE headset. The photodiode is attached to the micro-USB port of the MUSE and is used to provide a marker signal, which can be used to unambiguously align the recordings from different MUSE headsets. A python library for having a robust and stable Bluetooth connection between RPi and MUSE was developed and tested in naturalistic conditions in a cinema. The results show that this system proves to be resilient in terms of stability and data loss as well as accurate in temporal alignment between all headsets with the use of the photodiode. These two crucial characteristics, combined with its little necessary preparation time,. low cost and off-the-shelve availability confirm its suitability for a scalable hyper-scanning system.

## MATERIALS AND METHODS

### MUSE-RPi Coupled System Development

One of the primary specifications of the project was to develop a scalable system for deployment in a large audience or group. It was decided to purchase 10 MUSE headsets (the first 2016 version of the headsets) for developing a proof-of-concept such system.

### Initial Attempt with Existing Software

The first approach was to try to record all 10 MUSE headsets by a single computer. To test this option we used the proprietary software provided by the manufacturer of MUSE for connecting and recording data from it (InteraXon, 2022). The test was performed on a Windows 10 laptop, with a build-in Bluetooth adaptor. We progressively connected 4 MUSE devices. When only one headset was connected, a test recording was performed for 5 minutes, in which no data got dropped. Then a second MUSE was connected and recorded. In a 5 minutes recording, about 1 minute of data was dropped for both MUSEs. When the third MUSE was also connected and 5 minutes were recorded, for two of the MUSEs there was about 2 minutes of data dropped and for the third most of the data was dropped. With the fourth MUSE also connected, in a 5 minute recording, for two MUSES about 3 minutes of data was dropped and for the other two no data was recorded. This test showed that recording multiple MUSEs on a single computer with the proprietary software was not a feasible option.

The next option that was investigated was to develop a solution in which each MUSE would be recorded on a single computer. Of course, as one the main aims was to have a scalable system, such a computer should be as low-cost as possible, so that it is feasible to scale recordings to 10, 20 or 100 people without a prohibitively high cost. The cheapest identified solution identified at the time of the experiment was Raspberry Pi 3 B+(RPi-Foundation, 2022). It has an ARM Cortex-A53 processor at 1.4Gz, 1 GB of LPDDR2 SDRAM memory, build in dual band Wi-Fi at 2.4 and 5 Ghz and build-in Bluetooth 4.2/ BLE(Bluetooth Low Energy). These specifications are adequate for connecting to a MUSE headset through Bluetooth, receive data from it and record it on a file. Also, the low power consumption by the CPU offers the possibility to provide power through an external battery, making the system wireless and easier to deploy. Based on this evaluation 10 Raspberry Pi (RPi) computers purchased accompanied by 10 light-weight lithium batteries for powering them. A single RPi with attached battery is shown in Fig. 1B.

In terms of operating system, the Raspberry Pi runs a version of Debian Linux called Raspberry Pi Operating System(OS), which is open source(RPi-Foundation, 2022). There is no proprietary software provided by MUSE’s manufacturer for connecting and recording MUSE headsets on Linux computers. One of the most popular libraries for recording MUSE on linux is called MUSE-LSL(Barachant et al., 2019). This library is using as backbone the Lab Streaming Layer (LSL) library (LSL, 2022), which has been developed to standardize and unify synchronous data collection from various types of devices. MUSE-LSL has been developed for both windows and Linux and attempts have been done before to record data on Raspberry Pi with it. We followed one of these attempts, described in the blog of ADAFRUIT (ADAFRUIT, 2018). For connecting from a Linux computer to a Bluetooth device, MUSE-LSL is using the Python Module “pygatt” for Bluetooth LE Generic Attribute Profile(GATT). The MUSE headset is broadcasting the recorded data through different Bluetooth “Services” channels or slots. Specifically for MUSE there are different services for different types of data. These are shown in Table 1.

**TABLE 1.** Bluetooth Services of MUSE headset with corresponding Service Universal Unique Identifiers (UUID). The computer dedicated for recording from a MUSE must subscribe to the corresponding Bluetooth services. Most of the services entail data transfer from the MUSE to the Computer. In the case of the “Control” service messages can also transferred in the opposite direction towards MUSE, such as commands to restart it for example.

| SERVICES | Service Universal Unique Identifier (UUID) |
| --- | --- |
| EEG channel TP9 | '273e0003-4c4d-454d-96be-f03bac821358' |
| EEG channel TP10 | '273e0006-4c4d-454d-96be-f03bac821358' |
| EEG channel AF7 | '273e0004-4c4d-454d-96be-f03bac821358' |
| EEG channel AF8 | '273e0005-4c4d-454d-96be-f03bac821358' |
| EEG channel Right AUX | '273e0007-4c4d-454d-96be-f03bac821358' |
| 3-axis Gyroscope | '273e0009-4c4d-454d-96be-f03bac821358' |
| 3-axis Accelerometer | '273e000a-4c4d-454d-96be-f03bac821358' |
| Telemetry | '273e000b-4c4d-454d-96be-f03bac821358' |
| Control | '273e0001-4c4d-454d-96be-f03bac821358' |

Each of these services is broadcasting regularly packets of data. For example, for any of the EEG channels service, each packet comprises of an array of 13 numbers. The first number is the packet index and the following 12 numbers are 12 subsequent EEG recorded samples of the specific channel. In MUSE-LSL under Linux the pygatt library subscribes to each of these services, and continuously monitors if there are new packets broadcasted in these services.

Once a new packet has been broadcasted from any of the subscribed services, MUSE-LSL calls a function (typically termed “callback” function) that performs some specific operation on the received data. In the case of an EEG channel service, it unpacks the received data and appends it in a buffer dedicated for holding the EEG data. In MUSE-LSL the typical functionality is that this data buffer is further streamed to a network port (through the LSL library), where it can be assessed by other software on the same computer or by other computers for real-time analysis. Or alternatively the data buffer can just be saved in a file without any LSL involvement. One of the useful functionalities that MUSELSL employs from the LSL library is the recording of event markers. These event markers can be for example triggers from a stimulus presentation software representing the time that a command was issued to the graphics card to display a specific stimulus. LSL core functionality is to try to estimate the asynchrony between the different sources of data. In the specific example the two sources of data are the EEG data stream from MUSE through Bluetooth and the event markers stream representing when visual stimuli were sent to the graphics card in order to be displayed on the screen. In order to analyze for example the Visual Evoked Potentials from visual stimuli, it is necessary to know for each stimulus the exact time delay between an EEG sample been recorded and received by MUSE-LSL and the marker been issued by the stimulus presentation software. The LSL library has been developed to provide estimates of such delays so that different data streams can be synchronized with as little jitter as possible. Of course, there are many sources of variability that cannot be accounted for by LSL and which can introduce significant temporal jitter. In the specific example of visual stimuli, although the stimulus presentation software might send the command to the graphics card to present a stimulus at a specific instance, delays in the graphics card or other parts of the computer, such as shared memory, can cause skipped frames so that stimulus is presented with some uncertainty after one, two or more frames. This jitter can be significant. For a sample on a typical screen with a refresh rate of 60 frames per second, jitter of 1,2, or 3 skipped frames is translated into temporal jitter of 16.6, 33.3 or 50 mseconds respectively. Such temporal uncertainty can impact severely the estimation of a Visual Evoked Potential. LSL cannot account for this type of jitter, at least not through its default functionality.

Another type of temporal jitter comes from the side of the MUSE EEG headset. The MUSE headset does not transmit any timestamp with its data. Instead, each data packet has as header an incremental index number, the same for all 12 EEG samples contained in the data packet. MUSE-LSL at the beginning of each recording makes an initial estimate of the average delay of Bluetooth communication between MUSE and the recording computer. In the website of the LSL library it is mentioned that this is determined with an accuracy of 2 msec for wireless connections(LIBLSL, 2022). This is the accuracy of the estimated time difference between the transmission of the packet by MUSE and its reception by the computer. Based on this, the LSL provides the timestamp of the recorded data with respect to the clock of the recording computer. In practice this timestamp corresponds to all the 12 data samples contained in one packet of EEG data,. MUSE-LSL just checks whether there is a gap in the sequence of received packets and accordingly extrapolates the initially estimated timestamp to the current data packet based on the sampling frequency. With this strategy, error in the initial estimate is transferred to the timestamps of later data samples. Additionally, the extrapolation of the timestamps from an initial delay estimate soles entirely on the assumption that the sampling frequency is known very accurately. As already mentioned, each MUSE headset, according to its official specifications has a sampling frequency of 256 Hz, or 256 samples per second. However, it is not uncommon in consumer grade hardware the sampling frequency to be slightly offset from the official specifications. For short recording times such deviations can be considered insignificant. For example if the sampling frequency of a MUSE headset is 256.1 Hz, 0.1 Hz faster than expected, then in a recording of 10 seconds there will be 1 additional unexpected sample, in 100 seconds 10 additional samples, and in 10000 seconds of recording 1000 additional samples, which correspond almost to 4 seconds offset. So, in longer recordings, deviations of the sampling frequency can have an impact. So, it is vital to have a very accurate estimate of the sampling frequency for accurately estimating and extrapolating timestamps. Once the sampling frequency of the headset has been accurately determined, then this offset can be accounted for.

LSL has the capacity to continuously estimate and update the delay between MUSE and the receiving computer in real time. However, this would mean that in given instances the time-step between two successive data points would be different than the sampling interval of the MUSE and this cannot be the case as the sampling frequency is fixed. This would only be useful for real-time analysis where the interest is focus on a small time-window.

Now, imagine a case when the recordings of 10 MUSE headsets, each recorded on a RPi, have to be synchronized. Then there should be an LSL marker stream in a network where all RPis are connected, from which the RPis will receive a common marker to be saved along with the MUSE data. This marker can be used to align the recordings from the different headsets. This of course requires that the time it took for the marker to reach each of the RPis is known with high accuracy. Over a Wi-Fi network where a large number of devices is connected this is not a trivial problem. This will also be demonstrated in the last part of the results.

The above extended argumentation had purpose of demonstrating that the use of LSL for recording MUSE data and aligning it to event markers is not perfect and there are still various sources of significant jitter and uncertainty to be accounted for. Despite these limitations we first attempted to use MUSE-LSL in Raspberry Pie to record the data from an EEG headset. We followed installation instructions for Raspberry Pi from the Adafruit blog (ADAFRUIT, 2018). The installation took many hours especially due to scikit-learn. There was also an issue with permissions (See issue 1 in (Muselsl-Issues, 2022)) which had to be tackled.

After the installation of MUSE-LSL, the first test was performed with only one MUSE-RPi set. A stream channel was established to the MUSE headset by running in a terminal of RPi the command “muselsl stream –address 00:55:DA:B3:77:CE”(The latter part is the Bluetooth adaptor ID of the specific MUSE).

We tested the stability of this command by running it 10 times. A stream was successfully established in 8 out of 10 times The other 2 times there was a timeout of trying to connect to MUSE headset after 5 seconds.

We then tested recording of a single MUSE for 4 different durations, namely 5, 30, 60 and 120 minutes. The longer durations are of interest because one of the aims of this project is to have a scalable system that can record audiences during performances, which can have durations of 2 hours. One of the main issues of interest in these tests was how MUSE-LSL would deal with the fact that the size of the buffer used to store the MUSE data is dynamic, increasing everytime a new parcel is received in PRi. For a 2-hour recording of 5 EEG channels with a sampling frequency of 256 Hz and each EEG sample being a 12-bit unsigned integer the resulting necessary buffer would be 13.824 MB. During recording this buffer is increasing on-the-fly every time new EEG parcels arrive. This means, the RPi has to not only deal with subscribing to the Blueooth services of MUSE and getting each data packet, but also has to deal with increasing the data buffer size in memory every time a parcel is received. The second aspect is that MUSE-LSL saves the data buffer only at the very end of the recording. So if any issue occurs, the entire data in the buffer can be lost. So the single MUSE test was performed to test both the stability of the connection and the stability of Raspberry Pi performance. During these tests, all data was recorded successfully without and data parcels dropped.

The second test we performed was with all 10 MUSE headsets turned on at the same time, with each one been recorded on a different Raspberry Pi. The MUSE-RPi sets were placed along a single line, in close proximity with an average distance of 40 cm between them, resembling typical distances in an audience. This arrangement is depicted in Fig. 1G. The RPis were connected to a dedicated Wi-Fi network. A monitoring/control laptop, connected to the same network, was used to connect to each RPi through Secure Shell(SSH) protocol and start the MUSE-LSL process. This test recording was planned to last 20 minutes.

During this test serious problems were observed with connectivity between Raspberry Pi and MUSE headsets. The Bluetooth connection was repeatedly lost in all pairs resulting in extensive data loss. The test was restarted multiple times with the same connectivity issues occurring. Based on the error messages when the Bluetooth connectivity was lost the problem was identified to be with pygatt, the python wrapper for the linux library gatttool used for communicating and exchanging data with Bluetooth devices.

The problem appeared to be related to the close proximity of the Bluetooth devices. As close proximity was one of the primary specifications of the project, other options were investigated.

### Development from first principles

Based on the experience gained from studying and testing MUSE-LSL, we decided to develop from first principles and test a python library for recording multiple MUSE headsets in close proximity. The main aim of this system is to be deployed in typical auditoriums, where people are seated in close proximity. Emphasis was put on two parameters.

The first parameter of emphasis was stability in recording through Bluetooth under the presence of many devices in close proximity. To achieve this stability the following specifications were set:

As the recording is only to be performed on Raspberry Pi, a Bluetooth communication library was selected that was developed primarily for RPi. This library is Bluepy, a python interface to Bluetooth LE (Low Energy), primarily developed and tested on Raspberry Pi (BLUEPY, 2022).
Data is only recorded and stored in ASCII files on disk. No real-time processing is performed, in order to reduce the number of processes running during the data recording and potential bottlenecks that could hinder the acquisition of the data from the MUSE headset.
The data is not stored in a single buffer of dynamic size in memory for the entire duration of the recording, stored in a file on disk only at the very end of the process. Instead, the buffer in memory has a fixed, relatively small size and when it is filled, its content is written in a file on disk and new data starts filling it in. The fixed size of the buffer should be big enough to avoid very frequent writing to disk. This approach safeguards against loss of large amount of data in the case that the recording process crashes during the recording.

The second parameter of emphasis is the alignment of the recorded datasets from the different MUSE headsets with hardware markers of unambiguous timing. Although LSL was considered for broadcasting a stream of markers over Wi-Fi to all RPis, it was concluded that due to the various sources of potential jitter, discussed earlier, a much more robust solution would be to use a hardware marker, unambiguous for all MUSE-RPi sets and immune to any network instabilities.

#### Using a photodiode

The MUSE headset has an unused fifth auxiliary channel in the amplifier that samples the 4 EEG electrodes with a sampling rate of 256Hz. This channel is accessible through pin 4 of the micro-USB port of the MUSE headset. The electrode layout of MUSE and the micro-USB port are depicted in Fig. 1A. The reference electrodes for the EEG channels are on the central forehead. So, if a single EEG electrode is attached on pin 4 of the micro-USB then it is also referenced with the same reference signal. This is useful if the task at hand is to add an additional EEG electrode on a part of the head far from the default electrodes of MUSE EEG, such as visual or sensorimotor areas. In our case the task at hand is to add a sensor that will provide an accurate marker signal for aligning all MUSE headsets. The sensor used here was a photodiode. It provides a voltage across its two terminals, proportional to the intensity of the incident light. The specific photodiode was a planar photodiode, filter-coated for approximating the response of the human eye (LaserComponents, 2022) and is shown in Fig. 1C. The anode of the photodiode was connected to pin 4 of the micro-USB port and the cathode was connected to pin 5, which is the input ground pin of the micro-USB port. The photodiode wires were twisted and wrapped with conductive aluminum tape in order to minimize the contamination of the 4 EEG channels by the electromagnetic field created by the current flowing though the photodiode and its wiring (see Fig. 1D). The two main aspects of investigation were first whether this simple way of connecting the photodiode provides adequate voltage to capture light variations in experimental conditions and second whether there was any contamination from the photodiode channel into the other 4 EEG channels. These two aspects are both examined in detail.

#### RMPIMUSE Library development based on BLUEPY

BLUEPY python library has been developed primarily on RPi for communicating with Bluetooth Low Energy devices. Internally it employs code from the BlueZ project, the official Linux Bluetooth protocol stack(BlueZ, 2022).

Based on BLUEPY we developed a python library for recording data from a Muse headset in a paired RPi. The library is termed RPIMUSE in the rest of the manuscript and is available in GITLAB repository(RPIMUSE, 2022).

Apart from BLUEPY, many of the ideas used in the development of this library were inspired from some of the structure of MUSE-LSL, which was examined in the initial attempt to record the MUSE headsets through it.

In RPIMUSE the core class that handles most of data reception, unpacking and saving is RPIDelegate (located in rpimuse_delegate.py), which is inheriting the basic properties and methods from BLUEPY’s fundamental “Delegate” class. The higher level class that is responsible for all the main operations such as connecting and disconnecting to MUSE headset, and subscribing and unsubscribing to its Bluetooth services is the RPImuse class (located in rpimuse.py). This class uses BLUEPY’s Periperheral() class, with all the necessary functionality for connecting to MUSE Bluetooth services.

Some very basic parameters necessary for the above classes are stored and read from a script called rpimuse_constants.py. The main script that creates all the necessary objects and oversees the whole data acquisition process is run_rpimuse_inputs_reconnect1.py. This is the script that is called on command line with some input arguments and starts the acquisition. As input arguments one must set the hardware MAC address of the MUSE headset from which data is to be acquired, the duration of the recording, the directory where the data files will be stored and a mnemonic which will be used in the naming of the datafiles.

The MUSE headset has different sampling frequencies for its EEG and for its accelerometer and gyroscope devices. The EEG is sampled at 256 Hz while the accelerometer and the gyroscope at 52 Hz. As mentioned earlier, the data of each EEG channel is broadcasted through a different Bluetooth service. For the 3-axis Gyroscope this is not the case and the data for all its 3 axes are broadcasted through one service. The same happens with the 3-axis accelerometer. Regarding the size of each broadcasted data packet, things are also different for EEG and gyroscope/accelerometer. As already mentioned earlier the data packet for each EEG channel contains 12 consecutive data samples preceded by a single incremental packet number. The broadcasting of EEG data is performed in a cycle, where a data packet with a specific packet number is broadcasted in the channel sequence Auxilliary, TP10, AF8, AF7, TP9. So for the 5 EEG channels alltogether there are 5×12 data samples for the same packet index. Then the next data packet is broadcasted in the same sequence. This sequence is depicted in Table 2. It was decided to use for EEG a fixed size buffer corresponding to 30-second-long recordings. With EEG sampling frequency of 256 Hz the buffer size was equal to 30*256= 7680 samples for each channel, which fitted exactly 640 different data packets. As there are 5 EEG channels the total size of the buffer was 5×7680. Once this buffer was filled, the data was all written in an ascii file, the buffer was cleared and started getting filled anew. The datafile included in its name a file counter so that the data could be easily post-hoc concatenated.

**TABLE 2.**
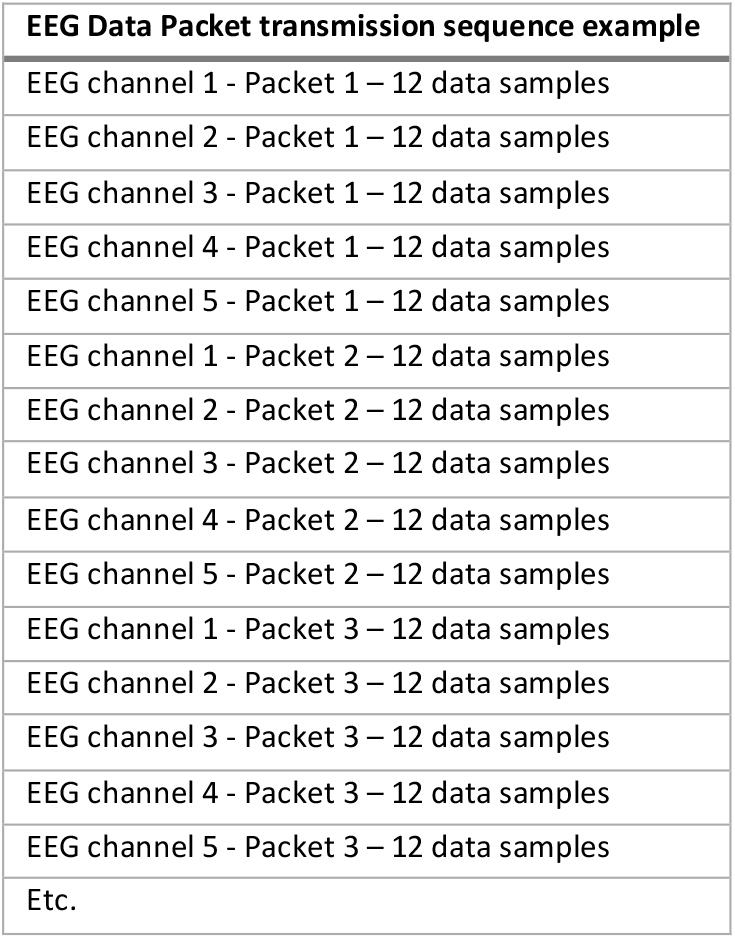
Sequence of transmission of EEG data packets from MUSE HEADSETS. Example of how data packets are broadcasted for the different EEG channels (4 default channels + Auxiliary). The transmission cycles through the different EEG channels for the transmission of recorded data corresponding to a time interval of 12 recorded consecutive samples.

| EEG Data Packet transmission sequence example |
| --- |
| EEG channel 1 - Packet 1 – 12 data samples |
| EEG channel 2 - Packet 1 – 12 data samples |
| EEG channel 3 - Packet 1 – 12 data samples |
| EEG channel 4 - Packet 1 – 12 data samples |
| EEG channel 5 - Packet 1 – 12 data samples |
| EEG channel 1 - Packet 2 – 12 data samples |
| EEG channel 2 - Packet 2 – 12 data samples |
| EEG channel 3 - Packet 2 – 12 data samples |
| EEG channel 4 - Packet 2 – 12 data samples |
| EEG channel 5 - Packet 2 – 12 data samples |
| EEG channel 1 - Packet 3 – 12 data samples |
| EEG channel 2 - Packet 3 – 12 data samples |
| EEG channel 3 - Packet 3 – 12 data samples |
| EEG channel 4 - Packet 3 – 12 data samples |
| EEG channel 5 - Packet 3 – 12 data samples |
| Etc. |

For the gyroscope the sampling frequency is 52Hz. Each data packet contains 10 unsigned integers with 16-bits resolution. The first number is the incremental index of the data packet from the beginning of the recording. The next 3 numbers represent one sample set, with one concurrent sample from each of the 3 axes of the gyroscope. The next 3 numbers represent the next sample set and last 3 numbers a third sample set. So each gyroscope data packet contains 3 samples for each axis. In this case the fixed buffer was sized so that it contains the same number of data packets as for the EEG, 640 data packets. As each data packet contains 3 samples of the 3-axes the total number of samples in each datafile is 1920 samples for each of the 3 axes. The sampling frequency, data packet size and fixed buffer size are identical for the 3-axis accelerometer.

Once a Bluetooth connection is established between the MUSE headset and the RPi, the RPi subscribes to the data Bluetooth services, the data packets start being broadcasted by MUSE and received by the RPi. As they are received, they are unpacked and their data is placed in the corresponding fixed size buffers. Once the buffers are filled, their content is written in a file on disk, then they are emptied and new packets are placed in them. When the recordings end, any content in the buffers is written in a last data file.

The above recording procedure is rather basic, performing only essential functions and avoiding any additional processes that could create bottlenecks in CPU or memory usage.

### TESTING

Testing was performed in two stages, one for testing the photodiode with a single MUSE-RPi set and one for testing all 10 RPi MUSES together for accessing the stability and robustness of this system.

#### Testing the Photodiode

In the first stage the suitability of the photodiode was examined as a source of a marker signal. This test was performed with a single MUSE-PRi set on a single participant, the experimenter.

The first test we performed was to investigate whether the photodiode produces voltage that could be used as a marker for inter-device alignment. The first level of this test was investigating the recorded signal in the auxiliary channel of the MUSE headset when the photodiode was placed on a monitor where a square was alternating between white and black color in equal intervals in a pulse-like fashion. We investigated different pulse durations in order to identify durations best suited for achieving a clean marker signal. Too short pulses could lead to a less well-defined marker due to the onset and offset slopes of the photodiode while too-long pulses can be corrupted by the high-pass filtering of the MUSE headset. This high-pass filtering is performed online by the MUSE headset because it has only 12-bit resolution and consequently slow voltage drifts could easily saturate the EEG channels. Thus real-time highpass filtering is necessary to avoid such situations. Typically, such high-pass filtering uses a cutoff frequency of 0.5 or 1 Hz which means that pulses with similar frequencies would be attenuated by the filters. We tested 8 different pulse lengths, namely 100, 166.6, 200, 400, 600, 800, 1000 and 2000 msec. During this test the MUSE headset was not placed on a participant.

#### Testing Contamination of the EEG channels from the Photodiode Signal

The next necessary step was to investigate to what extend the electromagnetic field produced by the photodiode current contaminates the other 4 EEG electrodes which are recording brain signal. The specific photodiode (LaserComponents, 2022) used in this study produces a maximum voltage of about 300 mV when connected to the auxiliary channel of MUSE headset and exposed to high luminance light. This voltage is at least an order of magnitude higher than typical EEG voltage levels. The photodiode is connected to the micro-USB part on the right posterior part of MUSE headset, very near the electrode T10, which runs behind the right ear (see Fig. 1A). Although the photodiode wires were twisted and shielded with aluminum conductive tape, still the connection to the micro-USB port is uninsulated. So, the current that runs through the photodiode wires creates an electromagnetic field which can contaminate the 4 EEG channels especially as the voltage from the brain is in the μV range. Electrode T10 is the most obvious candidate for being contaminated due to the very close proximity, but potentially all channels could be contaminated. By contamination it is meant that the photodiode signal will synchronously appear in the EEG channels, with reduced amplitude. A test recording was performed in order to investigate this possibility.

In this test the photodiode was attached to a piece of polyethylene terephthalate foam and taped on the shoulder of the participant (see Fig. 1E). The participant starred at a monitor 70 cm away from him, where a square of side length of 4 visual angle degrees was presented in the center (Fig. 1F). The square was alternating between black and white color every 200 msec, giving a total pulse length of 400 msec. There were 300 such pulses. Although a much smaller number of pulses would suffice to investigate the contamination of the electrodes from the photodiode current, it was decided to have a large number of pulses in order to investigate at the same time whether a visual evoked potential can be recorded by the EEG channels. So, in that context the 300 pulses corresponded to 300 trials. The signal from the auxiliary channel, where the photodiode was recorded, was first low-passed filtered at 15Hz, for removing any high frequency fluctuations.

#### Testing Group Recordings

The first test of recording of all the 10 MUSE-RPis together was performed in the lab with the headsets aligned in a line on the floor with an average distance of about 40 cm between them similar to the arrangement depicted in Fig. 1G. The RPis were connected to a dedicated Wi-Fi router. A monitoring/control laptop was also connected to the same Wi-Fi network. The rpimuse library was started through SSH in each RPi and then all MUSE headsets were turned on. The recording lasted for 30 minutes in each MUSE. The recorded files were then loaded and concatenated in analysis computer in MATLAB. The data packet indices, recorded in the data files, were examined in order to examine whether there was any loss of data. There was no EEG, Gyroscope or Accelerometer data loss at all in any of the MUSE headsets. This preliminary successful test provided an initial verification of the stability and robustness of the RPIMUSE library even when the 10 MUSE-RPi sets were in close proximity. The next step was to test the system with long recordings in naturalistic conditions, in order have a higher level of confidence in its stability.

A suitable opportunity for performing such a test was provided by a series of 4 events, termed together as “The Brain on Screen”, organized by the Max Planck Institute for Empirical Aesthetics in the German File Museum in Frankfurt, Germany. In each of these events a movie was played in the cinema of the German Film Museum (Fig. 2A), preceded by a talk from a neuroscientist. The name of the 4 movies played in these events are shown, along with their durations, in Table 3.

**Figure 2.**
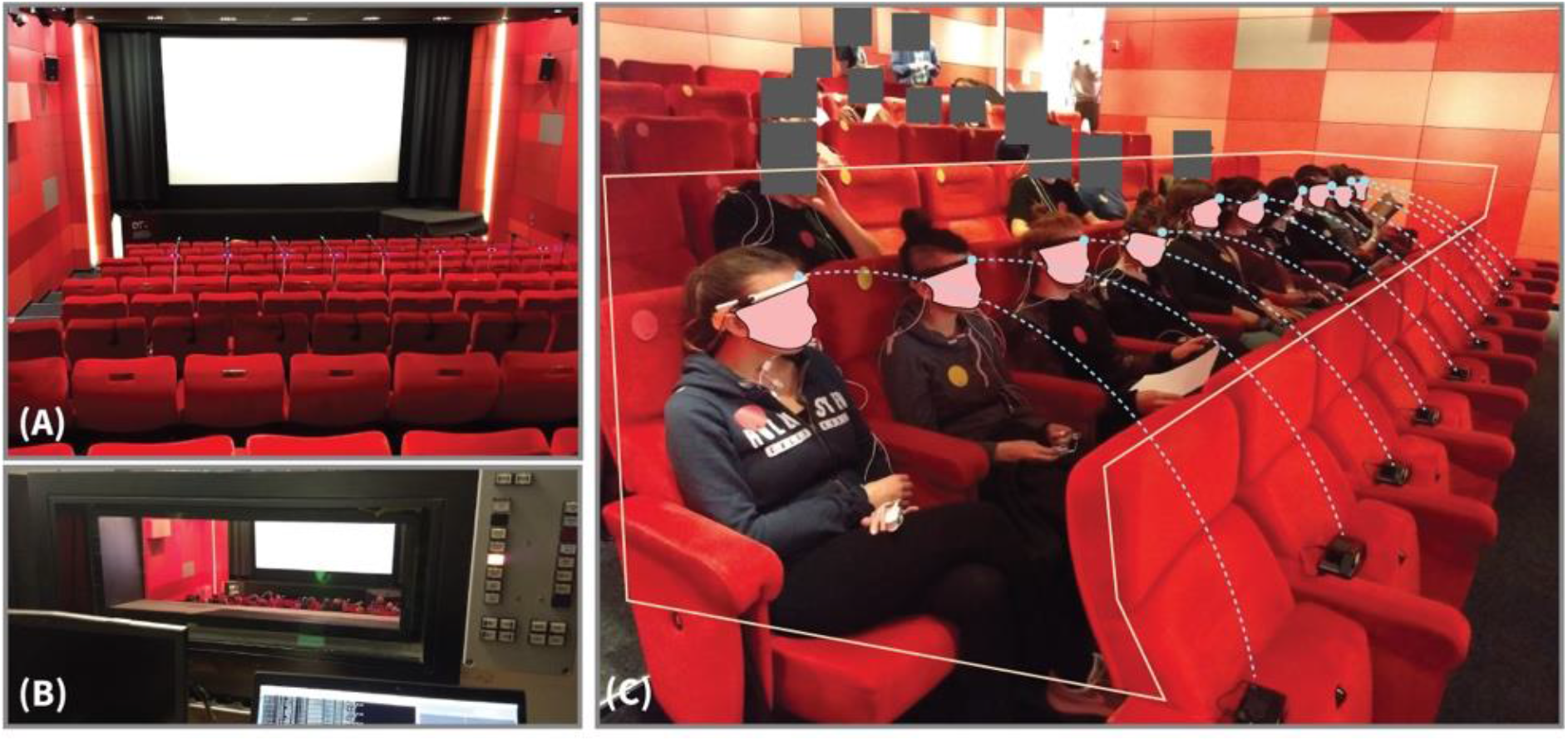
Setup of Multi-MUSE recordings in the German Film Museum. (A) The recordings took place in 4 different days during the showing of 4 different movies inside the cinema of the German Film Museum. A dedicated wi-fi router was placed inside the cinema for communication between a monitoring/control laptop computer and each of the 10 Raspberry Pi s. (B) The monitoring/control laptop computer was placed inside the projection room of the cinema. The monitoring of the recording and the communication with the Raspberry Pi s was performed entire from the projection room without any physical presence inside the cinema room. (C) The 10 participants were seated next to each other in a row (thin frame). The MUSE headsets were placed on their heads. Each one was connected to and recorded by a dedicated Raspberry Pi, placed on the seat right in front of the participant (pairing of MUSE headsets with dedicated Raspberry Pi s is depicted by the doted arcs).

**TABLE 3.** Movies played in the German Film Museum in the 4 events of the “Brain on Screen” project

|  |  |  |
| --- | --- | --- |
| **Movie 1** | Fear Eats the Soul (R.W.Fassbinder, 1974) | Duration: 89 min |
| **Movie 2** | Harold and Maude (H. Asby, 1971) | Duration: 91 min |
| **Movie 3** | After Hours (M. Scorsese, 1985) | Duration: 97 min |
| **Movie 4** | Run Lola Run (T. Tykwer, 1998) | Duration: 80 min |

The audience comprised of 100 people. The main overarching aim of this project was to perform a feasibility study of running Audience experiments in naturalistic settings. Under this aim, 55 members of the audience were the same across the 4 events and were seated in the same seats. The autonomic nervous system of each of these 55 participants was recorded using a Biosignals Plux system (PLUX, 2022) equipped with various peripheral sensors. A sub-group of 10 of these participants was selected for testing group EEG recording using the MUSE-PRi sets and RPIMUSE library. All 10 MUSE-RPi sets were equipped with a photodiode so that their data could be aligned post hoc.

A dedicated Wi-Fi router was placed inside the cinema room and a dedicated network was setup. In the router setup, a specific permanent IP address was assigned to each RPi, according to its Media Access Control (MAC) address. A monitoring/control laptop was placed inside the movie projection room at the rear of the cinema (Fig. 2B). This monitoring laptop was also connected to the dedicated Wi-Fi network and was used to first check that all 10 RPis were connected to the Wi-Fi router and also to connect through SSH to each one of them and start the RPIMUSE recording software. Although this way of starting the recordings was not the most sophisticated, it was chosen because it was the most direct way to see whether there were any issues in starting the recordings.

The 10 participants were all seated in the same row (see Fig. 2C). While they were still not in the cinema, the RPis were placed on the seats right in front of the seats where the corresponding MUSE headsets would be recorded. Each MUSE headset was placed on the back of the chair, where its corresponding participant would sit and it remained turned off. The RPis were then turned on by connecting their batteries (there is no turn-on button in RPIs). Once they were all connected to the dedicated Wi-Fi router, then through ssh connection from the monitoring laptop the RPIMUSE library was started for recording from the MUSE headsets. When RPIMUSE started in a specific RPi, it first searched for its dedicated MUSE headset and it remained in this state until it could find it. Once the RPIMUSE process was started in all RPis, slightly before the 10 participants entered the cinema, the 10 MUSE headsets were turned on while still on the back of the chairs. The RPIMUSE process running on each RPi was then connected to its dedicated MUSE headset and the data recording started. This was the beginning of MUSE data collection. Shortly after, the participants entered the cinema room and they sat on their seats. Then one experimenter helped each one of them to put the MUSE headset on and to make sure that the EEG sensors were attached on the head skin as much as possible. The participants remained in their chair until the beginning of the event.

The first part of the event inside the cinema room was a talk by a neuroscientist. Then there was a sequence of on-screen instructions for specific tasks such as eye-movements, turning the head left or right, breathing and other such movements, mostly for deriving templates of artifacts useful for the analysis of the peripheral measurements. After this sequence of tasks the movie was started. Regarding the EEG recordings, there was a sequence of 3 visual pulses on the screen just before the start of the movie and another identical pulse sequence right after the end of the movie. These visual pulse sequences were used as the markers picked by the photodiodes in order to align the recordings of the different MUSE headsets. After the end of the movie the participants were instructed, by an on-screen message, to remain to their seats and after 5 minutes the lights were turned on and the audience could leave the cinema room. The 10 participants with the MUSE headsets had been instructed to remain in their seats until an experimenter would come and remove the headsets from their heads. Before removing the headsets, the RPIMUSE process in each RPi was stopped through SSH from the monitoring laptop.

#### TESTING THE DELAYS OVER THE WI-FI NETWORK

During the recordings in the German Film Museum another small test was performed in order to test the stability of communication between the monitoring laptop and each of the MUSE headsets over the Wi-fi connection. This is relevant in order to get a first idea of whether data alignment over Wi-Fi could be feasible. We used one of the most popular open-source universal messaging libraries called ZEROMQ (ZEROMQ, 2022). The monitoring/control laptop ran a python script during the movie recordings, which opened a TCP socket in the dedicated Wi-Fi network and broadcasting to the network its timestamp every 0.5 seconds. In each RPi a small function, added to the RPIMUSE library, was checking in the specific TCP network socket whether a message had been broadcasted. Once a new message was broadcasted the RPi muse recorded it into disk after adding next to it its own timestamp of the moment the message was recorded. The difference between the timestamp contained in the broadcasted message and the timestamp of the time it was received by the RPi represented the communication delay over the Wi-Fi network.

### ETHICS STATEMENT

The experiments were approved by the Ethics Council of the Max Planck Society. Written informed consent was given by all participants before the experiment.

### DATA ANALYSIS

The analysis of the recorded EEG and photodiode signals was performed with MATLAB (MATLAB, 2022) and with Fieldtrip toolbox (Oostenveld et al., 2011).

## RESULTS

### TESTING THE PHOTODIODE

We first tested with different visual pulse durations on the monitor the signal recorded by the photodiode in MUSE. We tested 8 different pulse lengths, namely 100, 166.6, 200, 400, 600, 800, 1000 and 2000 msec.

Fig. 3A presents a segment of the recorded photodiode signal in the auxiliary channel for a visual pulse duration of 400 msec. The onset of the visual pulses is clearly visible at the very beginning of the interval and thereafter the photodiode signal appears to track well the pulse sequence. This verifies the validity of the photodiode signal as a potential marker. In order to quantify the identification of a pulse pattern in the photodiode signal, it was convoluted with a pulse kernel. This kernel is depicted in Fig. 3B for a pulse length of 400 msec band a total duration of 5 pulses. This kernel slides through the photodiode signal and its convolution is computed at every timepoint. Fig. 3C to 3J depict 2 subsequent recorded photodiode pulses for different pulse lengths (upper subplot) and the corresponding convolution curves (bottom subplots). For the shortest pulse duration of 100 msec (Fig. 3C) the photodiode signal seems to not be able to capture well the pulse shape. As the pulse length increases, the photodiode signal captures progressively better the shape of the pulse up to a pulse length of 400 msec. Thereafter for longer pulse lengths there is the introduction of progressively longer additional noise towards the end of the down and upper phases of each pulse, with the pulse shape being significantly distorted for the longest durations of 1000 and 2000 msec. This additional noise is most likely introduced by the real-time high-pass filtering of the MUSE headset and that is why is manifested only in the longer pulse durations. There the pulses are cut short by the filter which tries to eliminate low frequencies. An additional source of the noise in the down phase of each pulse comes from the electromagnetic noise of the environment picked up by the photodiode and its wiring. In the up phase of the pulse, when the on-screen rectangle is white, the photodiode produces current which creates the voltage difference across the two terminals of the photodiode. When the on-screen rectangle becomes black the photodiode stops producing current and the voltage returns to the down-phase baseline. However due to the electromagnetic noise of the environment, such as changing magnetic fields, eddy currents are introduced in the photodiode and its wiring, which can fluctuate randomly and which are translated into noise fluctuations during the down-phase of the photodiode pulse patterns. As mentioned earlier the photodiode wires were twisted and covered with aluminum conducting tape in order to reduce such noise. However, some noise still persisted and this is can be due to the least shielded parts of the circuit, the photodiode and the micro-USB connector. From the photodiode signal plots in Fig 3C to 3J (upper subplots) it can be concluded that a pulse duration around 400 msec seems be an optimal value where the shape of the pulse is captured well, while noise from the real-time filtering of the MUSE headset and the electromagnetic environment has a small effect. Of course, the electromagnetic noise depends on the given environment during a recording and these results presented here are just used in a proof-of-principle fashion rather as strict guidelines. In the bottom subplots of Fig. 3C to 3J are depicted the convolution curves of the photodiode signal with the 5-pulses-long convolution kernel. Apart from the shortest pulse duration of 100 msec (Fig. 3B), where the pulse shape is highly distorted, for all other pulse lengths the convolution curve has a peak (transparent red line), that corresponds to the sharp onset of the pulse. Even in the case of the longest durations of 1000 and 2000 msec, where the pulse shape is highly distorted by the high-pass filter of MUSE and noise, the convolution still has its maximum at the pulse onset. In conclusion, the convolution with a template pulse kernel seems to be a robust method, tolerant to high levels of noise for identifying the pulses in the recorded photodiode signal.

**Figure 3.**
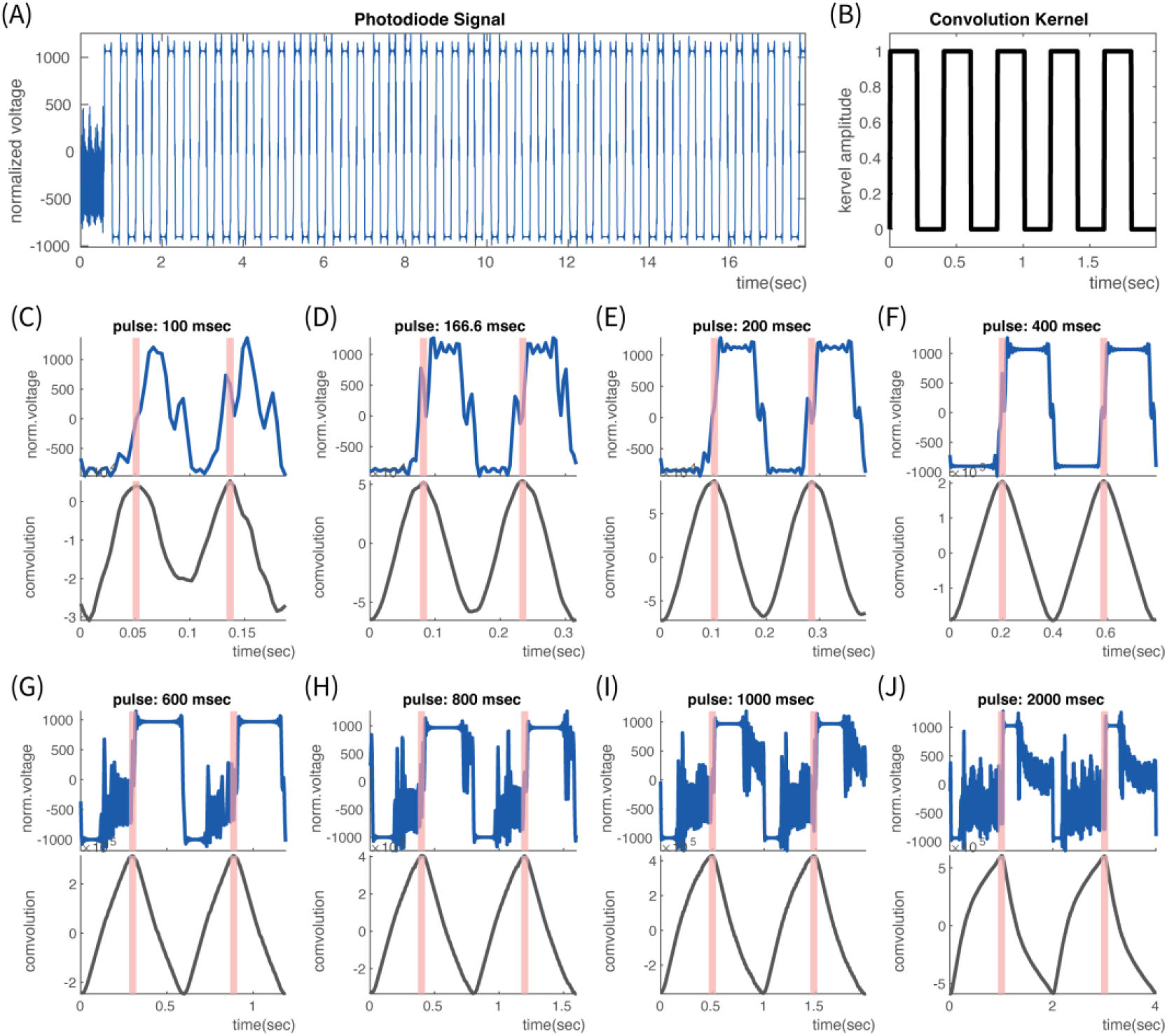
Off-head Photodiode Testing with different pulse durations. (A) Example of Photodiode time-series when the duration of the visual pulse is 400 msec (200msec on-200 msec off). At time zero the pulse sequence has not started yet. Shortly after the pulse sequence starts and the photodiode signal clearly captures the pulse sequence. (B) Convolution 5-pulse-long kernel. Here for a pulse length of 400 msec. This kernel is convolved with the photodiode signal in order to quantitively identify the onset of each pulse. (C) to (J) Representative segment of photodiode signal(upper subplot) and corresponding convolution with 5-pulse kernel (lower plot) for various pulse durations. The transparent red lines mark the convolution peaks and the corresponding marked onset points on the pulses in the photodiode signal. Pulse durations: (C) 100 msec, (D) 166.6 msec, (E) 200 msec, (F) 400 msec, (G) 600 msec, (H) 800 msec, (I) 1000 msec, (J) 2000 msec. Note that for the shortest durations in (C) and (D) the photodiode signal does not capture so well the pulse shape. This is better captured in the case of 200 and 400 msec pulse durations (E) and (F). For longer pulse durations, (G) to (J) noise is introduced in the down- and up-phase of the pulse due to the real-time high pass filtering in MUSE and due to noise in the environment.

### TESTING CONTAMINATION OF THE EEG CHANNELS FROM THE PHOTODIODE SIGNAL

Having established that the photodiode provides a signal that can be used as a marker, we then tested whether the electromagnetic field produced by the current in the photodiode contaminates the recordings of the other 4 EEG electrodes. As described in Methods, this was tested by an on-head test during which on the screen was presented a sequence of 300 visual pulses, each on with a length of 400 msec.

A characteristic segment of the photodiode signal is shown in Fig. 4A, where it is clear that the photodiode captures well the visual pulses, as expected from the off-head test with the same pulse duration already mentioned in Fig 3F. The main difference observed here in the photodiode signal in Fig. 4A is that there is more noise towards the end of the bottom-phase, just before the onset of each pulse. This is possible due to the environmental noise conditions during the on-head testing as compared to the off-head testing. Despite this increased noise, the convolution with a convolution kernel of length of 5 pulses provided a consistent pulse onset identification, as seen in Fig. 4B. The time instances of the convolution peaks are depicted by the transparent red lines.

**Figure 4.**
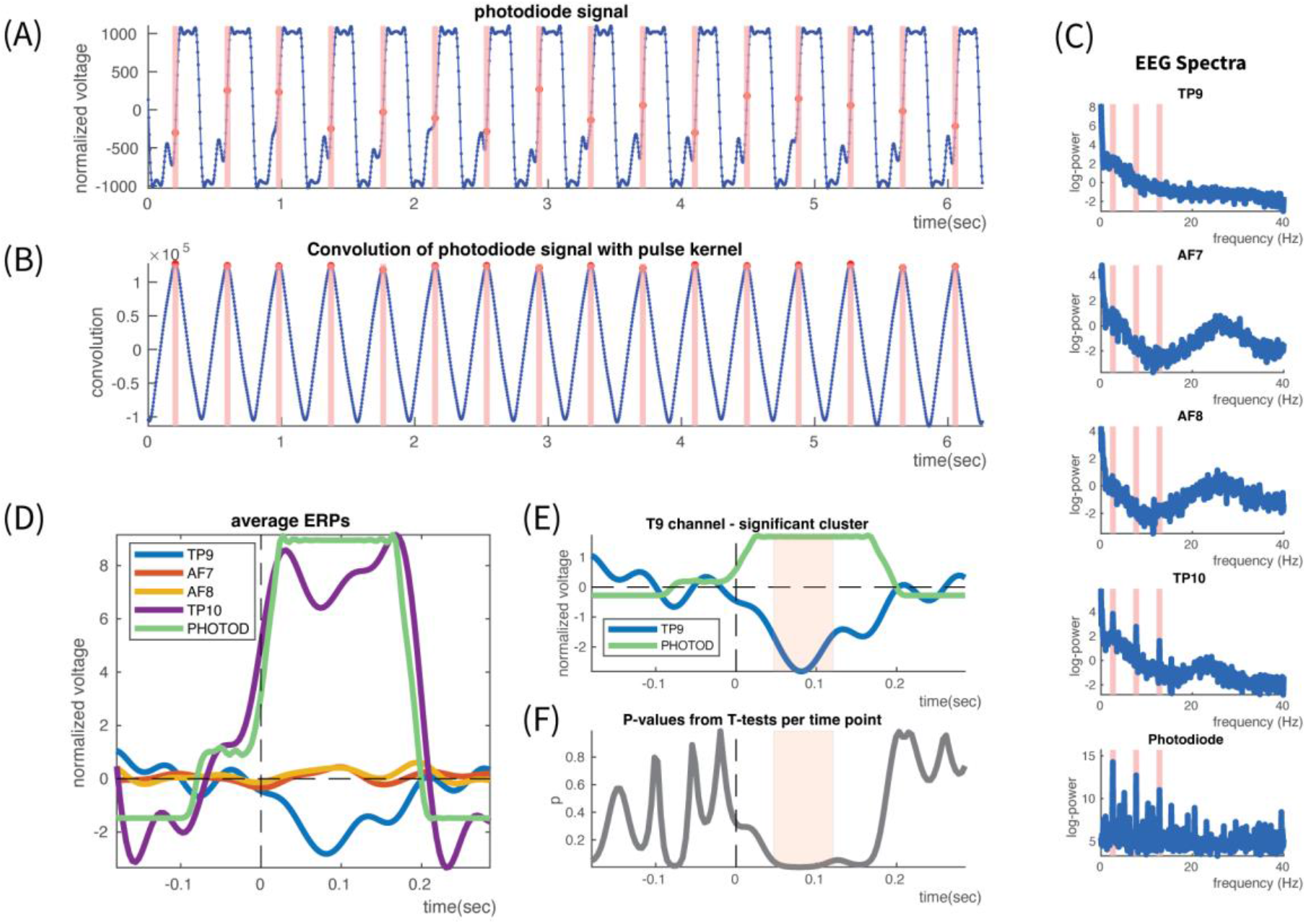
On-head Photodiode Testing. Performed with a visual pulse duration of 400 msec (200 msec on and 200 msec off). (A) Representative segment of the photodiode signal. The shapes of the pulses are clearly captured by the photodiode. Some noise is present in the later down-phase of the pulse just before onset. (B) Convolution with a 5-pulse convolution kernel. The clear peaks provided consistent pulse onset identification, marked by transparent red lines (which also extend above to show on the photodiode signal the identified onsets). (C) Log-power spectrum for each of the 4 electrode channels and the photodiode. As the expected, the Photodiode has the strongest power peak at 2.5 Hz (very narrowband, spike-like), which corresponds to the visual pulse duration of 400 msec. It also has prominent narrowband peaks in many higher harmonics. The two harmonics with the highest power are the 3^rd^ and 5^th^ at 7.5Hz and 12.5 Hz respectively. The pulse frequency and these 2 harmonics are marked with transparent red lines which extend above to the spectra of the EEG channels in order to investigate whether they are contaminated at these frequencies. Channel TP10, in close proximity to the micro-USB port, is clearly contaminated in these 3 frequencies, at it has corresponding narrowband peaks in its spectrum. The other channels are not contaminated in these frequencies. (D) This is also confirmed from the average ERP of the EEG and photodiode channels. Time 0 is the pulse onset identified in each pulse in the photodiode channel. The ERP of TP10 clearly tracks the average photodiode signal. The other EEG channels do not appear to be contaminated from the photodiode. The ERP of the left posterior electrode TP9 appears to have a visual evoked potential during the up-phase of the photodiode, peaking in the vicinity of 100msec. Interestingly also the contaminated channel TP9 seems to have the same visual evoked potential superimposed on the photodiode artifact. (E) Cluster-based permutation statistics showed that the peak of the ERP in channel TP9 is significant around 100 msec(red transparent area), hinting that this could be P100, with inverted polarity due to the lateral location of the sensor. (F) The P-values from the individual, per time-point, t-tests used in the cluster permutation statistics. Beyond the region around 100 msec which was marked as significant (red transparent area), it is also seen that in the vicinity of 170 msec there are also low p-values, although not surviving the statistical significance threshold. Another hint that this is probably visual evoked potential recorded in posterior sensors.

The next step was to compute the power spectrum of each EEG channel and the photodiode and see whether the frequency of the visual pulse is not only present in the latter but also in any of the former.

The spectrum was computed in Fieldtrip toolbox (Oostenveld et al., 2011) using a moving window with a Hanning taper at each time point of recorded data. The investigated frequencies were between 1 and 40 Hz with a step of 0.25 Hz. For each frequency the moving window had a different length, equal to 5 times the period of the investigated frequency. For example, for 20 Hz the moving window had a length of 5*1/20=0.25 sec while for the slower 10 Hz had a length of 5*1/10 = 0.5 sec. At each time point the moving windows for all the investigated frequencies were centered to it and the spectrum was computed. This was repeated for each time point providing a time-series of spectra. These spectra were then averaged across time and provided the average spectra for each EEG channel and the photodiode, which are presented in Fig. 4C. In the spectrum of the photodiode, at the very bottom, the frequency with the strongest power is 2.5 Hz, which corresponds to the frequency of the visual pulse ( ie. Pulse duration of 0.4 sec corresponds to 2.5 Hz frequency). The spectrum in this frequency is characteristic of an extremely regular pattern as it is very narrowband with a spike-like shape. Apart from the external stimulus frequency itself, there are similar peaks in the photodiode spectrum in the harmonics of this frequency. The harmonics with the next highest power are the 3^rd^ and 5^th^ harmonics, at 7.5Hz and 2.5Hz respectively. On Fig.4C these are marked with red transparent lines, which run also all the way up through the spectra of the EEG channels, in order to assist their examination for possible contamination from the photodiode current. Following these lines, it is easily seen that the right temporo-parietal electrode T10 has similar spike-like spectral peaks in the visual pulse frequency as well as the 3^rd^ and 5^th^ harmonics. T10 is the electrode in close proximity to the micro-USB port (Fig. 1A) and its spectrum confirms that the current flowing through the photodiode into the micro-USB port creates an electromagnetic field that contaminates it. The spectra of the other electrodes in Fig. 4C do not have any such peaks and thus they seem to have no contamination by the photodiode.

To further investigate the effect of the photodiode in the EEG recording we computed the average Evoked Response Potential (ERP), by epoching the recordings at the onset of each pulse, splitting accordingly the data into trials and averaging across them. With time 0 being the onset of each pulse, each trial spanned between −200 to 300 msec with respect to it. Each such trial was baselined based on the interval [-150 0] msec. The average ERP of each EEG channel and the photodiode are shown in Fig. 4D. The contamination of electrode T10 is here even more pronounced, as its ERP tracks in overall the ERP of the photodiode. The other electrodes do not show any such pattern and they seem to be unaffected by it.

The ERP of the left temporo-parietal electrode T9 during the up-phase of the pulse follows a transient pattern, similar to that of a visual evoked potential. It has two negative peaks just before 100 and 170 msec, resembling P100 and N170. The first peak here is negative but this possibly due to the fact that the T9 electrode is located relatively far from the occipital areas and a big part of it covers an area over the ventral part of the brain. There the P100 potential can be reversed. It has been shown that the P100 component originates in the bottom of the calcarine fissure and it reverses its polarity ventrally (Seki et al., 1996). This area of reversal includes largely the area behind the ear where the T9 electrode is located. So, the negative peak around 100 msec is possibly due to this reason. The fact that the peak occurs earlier than 100 msec is due to the method of pulse onset identification. As it can be seen on Fig. 4D, on the ERP of the photodiode, the pulse onset identified by the convolution method is marked on the rising slope of the photodiode onset rather at its very beginning. This can be possibly due to the noise present in the photodiode signal just before pulse onset, which is depicted as a little step just before zero. This positive offset in the identification of the pulse onset is the main reason that the first peak of the T9 ERP appears to occur before 100 msec.

The same is also the case for the second peak which appears to occur before 170. Based on these results and their interpretation, it appears that T9 captures the visual evoked potential from the visual pulse onset on the screen. Interestingly a visual comparison between the ERPs of T9 and T10 on Fig. 4D shows that the same pattern appears also in the ERP of the contaminated channel T10, superimposed on the artifact of the photodiode pulse. So, although channel T10 is contaminated by the photodiode signal it still captures brain activity. The validity of the visual evoked potential observed in the ERP of the T9 was statistically tested. Using Fieldtrip toolbox, we performed a non-parametric, cluster-based permutation statistical test (Maris & Oostenveld, 2007) in order to identify which parts of the ERP of T9 where statistically different than 0. The results are show in Fig. 4E, represented by the red shaded area. The first negative peak was found to be highly significant with p < 0.0099 and average cluster t-statistic=-101.32. The cluster extended in the interval [0.55 to 1.15] msec. The second peak just before 170 msec was not found to be significant according to the cluster statistics. By giving a look to the p values of each t-test, in each individual time point, in Fig. 4F, it can be seen that they are very low in the area of the significant cluster but also around the second peak before 170 msec. This means that although this second peak did not pass the statistical significance threshold, still a statistical trend is obvious there a fact that reinforces that the observed pattern is the visual evoked potential.

In summary, there were two main results. First, the electromagnetic field produced from the photodiode current contaminates electrode T10, right next to the micro-USB port, while the other 3 EEG channels are not affected. Second, after marking the onset of pulses by the photodiode signal, the average ERP of channel T9 captures the visual evoked potential from the pulse onset. The same ERP is also present in channel T10 superimposed on the photodiode artifact. These results show that although the photodiode contaminates one EEG channel, brain signals can still be successfully recorded by the MUSE in the other 3 EEG channels. Finally, even in the contaminated EEG channel, brain activity is captured (which could potentially be separated to some degree from the photodiode artifact if for example the variation of the light is known).

### TESTING GROUP RECORDING IN NATURALISTIC CONDITIONS

As described in Methods, the entire assembly of the 10 MUSE-RPi was put to the test in 4 long recordings during four events, including movie showing, in the cinema of the German Film Museum.

For all events, the time interval between the beginning of recording of data from each MUSE data in the corresponding RPi, just before the participants entered the cinema room, and the end of the recording, after the end of the movie, was in the order of 2.7 hours. Table 4 presents in detail the total recording time for each MUSE-RPi pair, in each of the four movies.

**TABLE 4.** Duration of recordings (in minutes) of each MUSE headset during each movie showing in the German Film Museum. All recordings were in the order of 2.7 hours. The recordings started much earlier than the start of the movie.

|  |  | MUSE HEADSET ID |  |  |  |  |  |  |  |  |  |
| --- | --- | --- | --- | --- | --- | --- | --- | --- | --- | --- | --- |
| MOVIE ID |  | 1 | 2 | 3 | 4 | 5 | 6 | 7 | 8 | 9 | 10 |
|  | 1 | 167.50 | 168.00 | 168.01 | 168.00 | 168.13 | 168.50 | 168.50 | 168.51 | 168.50 | 169.00 |
|  | 2 | 165.00 | 165.02 | 165.00 | 165.50 | 165.50 | 165.50 | 166.00 | 166.00 | 166.00 | 166.00 |
|  | 3 | 160.50 | 160.75 | 161.00 | 161.00 | 161.10 | 161.10 | 161.26 | 161.20 | 161.51 | 161.52 |
|  | 4 | 160.02 | 160.03 | 160.00 | 160.00 | 160.02 | 160.00 | 160.52 | 160.00 | 160.53 | 160.50 |

There were two primary aims of these long MUSE recordings. The first was to examine whether there is significant data loss when multiple MUSE headsets are recorded in close proximity under naturalistic noise conditions in an auditorium. The second aim was to examine whether the photodiode signal could be successfully used to align the recordings of the 10 different MUSE headsets in each movie under such naturalistic conditions.

Regarding dropped data, the recording of each MUSE for each movie was analyzed and the number of dropped data samples was computed from gaps in the sequential data parcel IDs that accompany each transmitted MUSE data packet (see Methods). The percentage of dropped samples for each MUSE in each movie is presented in Table 5.

**TABLE 5.** Percentage (%) of dropped data (EEG, Accelerometer and Gyroscope) for each MUSE headset during each movie showing in the German Film Museum. In the majority of cases there is no data drop at all. In all cases that there is data drop, it represents a very small percentage of the recorded data. Notice that the values are not ratios but already represent percentages (%).

|  |  | MUSE HEADSET ID |  |  |  |  |  |  |  |  |  |
| --- | --- | --- | --- | --- | --- | --- | --- | --- | --- | --- | --- |
| MOVIE ID |  | 1 | 2 | 3 | 4 | 5 | 6 | 7 | 8 | 9 | 10 |
|  | EEG |  |  |  |  |  |  |  |  |  |  |
|  | 1 | 0 | 0.002% | 0.008% | 0 | 0.079% | 0 | 0 | 0.004% | 0 | 0 |
|  | 2 | 0 | 0.015% | 0 | 0 | 0 | 0 | 0 | 0.002% | 0 | 0 |
|  | 3 | 0 | 0.157% | 0 | 0 | 0.063% | 0.061% | 0.161% | 0.125% | 0.004% | 0.010% |
|  | 4 | 0.015% | 0.021% | 0 | 0 | 0.012% | 0 | 0.010% | 0 | 0.017% | 0 |
|  | ACCELEROMETER |  |  |  |  |  |  |  |  |  |  |
|  | 1 | 0 | 0.005% | 0.009% | 0 | 0.163% | 0 | 0 | 0.002% | 0 | 0 |
|  | 2 | 0 | 0.009% | 0 | 0 | 0 | 0 | 0 | 0.002% | 0 | 0 |
|  | 3 | 0 | 0.271% | 0 | 0 | 0.037% | 0.057% | 0.222% | 0.216% | 0.007% | 0.017% |
|  | 4 | 0.019% | 0.033% | 0 | 0 | 0.019% | 0 | 0.014% | 0 | 0.019% | 0 |
|  | GYROSCOPE |  |  |  |  |  |  |  |  |  |  |
|  | 1 | 0 | 0 | 0.005% | 0 | 0.154% | 0 | 0 | 0 | 0 | 0 |
|  | 2 | 0 | 0.009% | 0 | 0 | 0 | 0 | 0 | 0 | 0 | 0 |
|  | 3 | 0 | 0.238% | 0 | 0 | 0.007% | 0.036% | 0.143% | 0.183% | 0.005% | 0.015% |
|  | 4 | 0.014% | 0.021% | 0 | 0 | 0.012% | 0 | 0.012% | 0 | 0.012% | 0 |

In the majority of cases there was no dropped data at all. In the cases when data was dropped, it always represented a very small percentage of the total recorded data. There was no MUSE headset that showed significant data loss in any of the recordings. The overall summarizing statistics of dropped data are shown in Table 6.

**TABLE 6.**
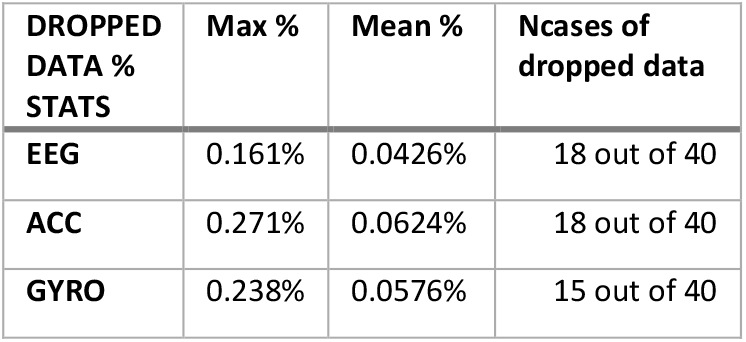
Summary statistics of the percentage (%) of dropped data (EEG, Accelerometer and Gyroscope) in the German Film Museum. The summary statistics confirm that the percentage of lost data is extremely small. Notice that the values are not ratios but already represent percentages (%).

| DROPPED DATA % STATS | Max % | Mean % | Ncases of dropped data |
| --- | --- | --- | --- |
| EEG | 0.161% | 0.0426% | 18 out of 40 |
| ACC | 0.271% | 0.0624% | 18 out of 40 |
| GYRO | 0.238% | 0.0576% | 15 out of 40 |

The EEG recordings had the smallest percentage of dropped samples with a mean of 0.0426% and a maximum of 0.161%. The gyroscope and the accelerometer had slightly higher percentages of dropped data but still in extremely low levels. These results confirmed that the developed RPIMUSE library, running on an RPi, can provide a very stable and consistent system for recording MUSE headsets, even if they are in close proximity in an auditorium. Another encouraging fact along this direction was that in the recording log files of each RPi, the of data loss were not accompanied by Bluetooth disconnection messages. Although it is not clear why these packages were dropped, one of the most probable reasons is that each RPi had an additional process running, which was monitoring the WI-FI network for possible control messages from the ZEROMQ stream of the monitoring laptop (see Methods). This additional process could be responsible for causing a CPU, memory or network adaptors’ bottleneck in the RPi at moments of data packet reception and this might caused the packet to be dropped.

The other aim of this experiment was to see if the photodiode can be used in a naturalistic audience environment for aligning the recordings of the 10 different MUSE headsets. As already described in Methods, just before the onset of each movie there was a sequence of 3 visual pulses presented on the screen of the cinema. An identical pulse sequence was presented just after the end of the movie. We used a 3-pulse kernel which was convolved through the recording of each MUSE in order to identify the onset of these pulses in the photodiode signal.

In Fig. 5A is shown the entire photodiode time-series from a single MUSE headset during Movie 1. Time 0 is the onset of the first marker pulse just before the start of the movie. This Start Marker Pulse can be clearly seen for Movie 1 in Fig. 5B, nicely aligned across all MUSES. The same consistent alignment was achieved in the other 3 movies as can be seen in Fig5C to E. As this Start Marker Pulse served as the reference time 0 and was consistently aligned across MUSES, it was then expected that also the first pulse after the end of the movie, termed End Marker Pulse, would be consistently aligned across all MUSE headsets’ recordings. This End Marker Pulse is shown in Fig. 5F for Movie 1. It is obvious that this pulse is not aligned across the different MUSE headsets. Instead, there seemed to be time differences between these pulses, in some cases in the order of 300 msec. Having made sure that this is not due to dropped data, the only cause of this misalignment could be that each MUSE headset had a slightly different sampling frequency in the vicinity of the official sampling rate of 256 Hz. This hypothesis was reinforced by the fact that the misalignment between different MUSE headsets was identical also in the recordings of the other 3 Movies in Fig. 5G to I. The next step was to estimate the corrected sampling rate for each MUSE headset. This was done using the recordings from Movie 1. MUSE headset 1 served as the reference headset with a reference sampling frequency of 256 Hz. Then for each MUSE headset, based on the time difference between the onset of the End Marker Pulse in its own recording and the onset of the same pulse in the reference MUSE headset, the new sampling frequency was estimated, which would eliminate this temporal offset. This was repeated for all MUSES. Based on the corrected sampling frequencies, the timestamps of all data points were recomputed. The resulting corrected End Marker Pulses for all MUSES are shown for Movie 1 in Fig 5J. As expected, the corrected sampling frequencies eliminated all temporal offsets. Using the already estimated new sampling frequencies, the timestamps of all data points were recomputed for all MUSE recordings in the other 3 movies. As it can be seen in Fig. 5K to M, the temporal offsets were completely removed and the End Marker Pulses were consistently aligned across all MUSES. This verified that the corrected sampling frequencies were valid.

**Figure 5.**
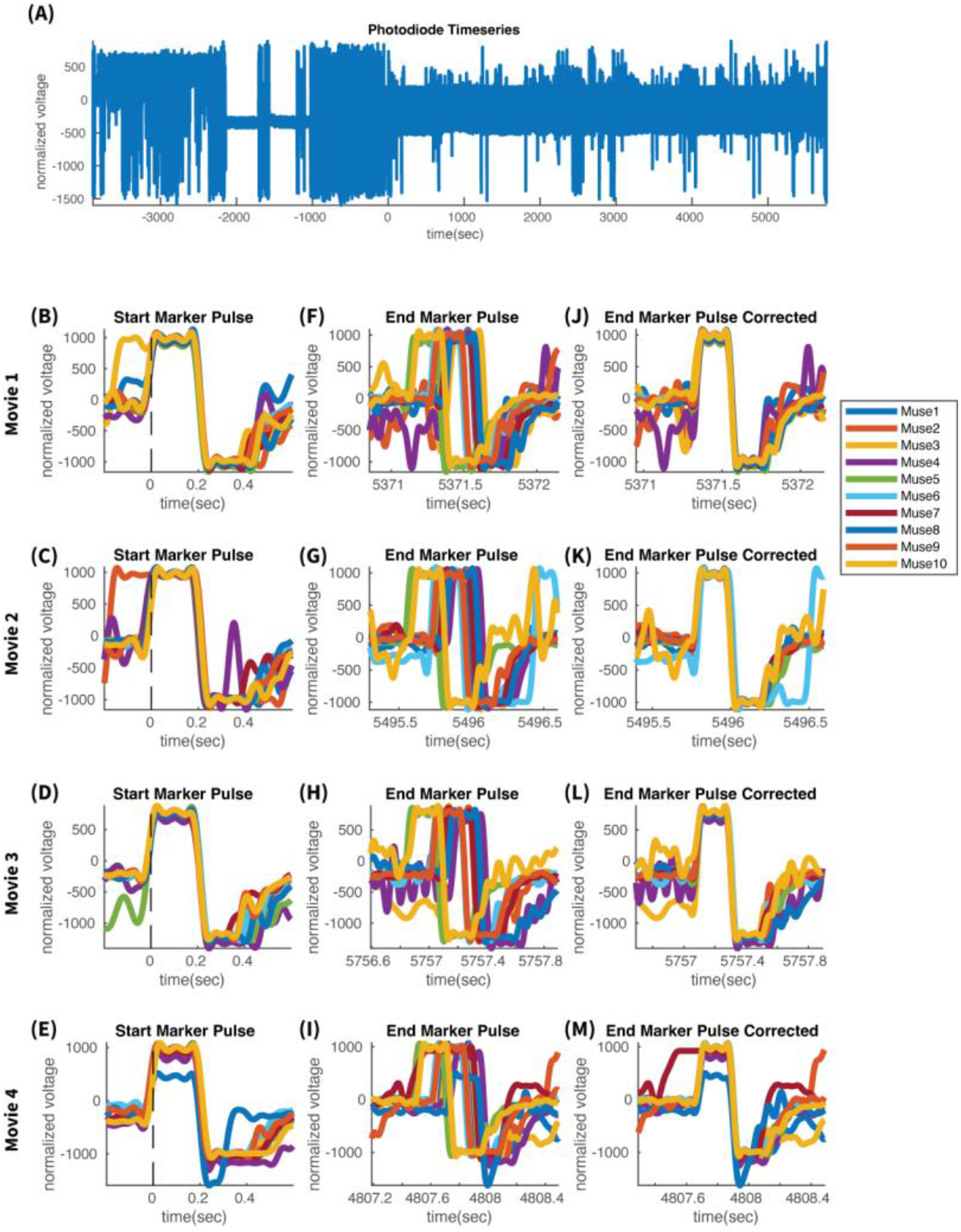
Alignment of Multi-MUSE data recordings. (A) The entire recorded photodiode time-series in Muse 1 during showing of Movie 1 in the German Film Museum. At time zero is onset of a visual pulse, used as a marker right before the beginning of the movie, termed here as Start Marker Pulse. A similar visual marker pulse occurs also after the end of the movie, termed here End Marker Pulse. Note that the recording starts much earlier than the beginning of the movie. The total duration of the recording is in the order of 2.7 hours for each of the 4 different movies. (B) In Movie 1 recordings the Start Marker Pulse is well aligned for all 10 Muse headsets, as the onset of this pulse was selected as the reference 0 time-point. The same happens in all the other 3 movies as depicted in (C) to (E). (F) The End Marker Pulse, after the end of Movie 1, is not well aligned across the 10 different MUSE headsets. For some MUSES, like 3(yellow) and 5(green) it occurs earlier while for other it occurs later. The onset time differences between MUSES do not exceed 0.4 seconds. After controlling for dropped data packets, the reason behind this misalignment was found to be the slightly different sampling frequencies of the different MUSE headsets. The same misalignment of End Marker Pulse was observed also in the other 3 Movies as depicted in (G) to (I). (J) In the recordings of Movie 1, when the timestamps of the data points were re-estimated with the corrected sampling frequencies (see Table XXX), the misalignment of End Marker Pulse was corrected and appeared well aligned across all headsets. With the same corrected sampling frequencies the End Marker Pulse was well aligned also in the recordings from the other 3 movies, as depicted in (K) to (M).

The corrected sampling frequencies are shown in Table 7. Their differences from the official sampling frequency of 256 appear very small for the purpose of a relatively short recording. However, for some headsets the sampling frequency difference can cause significant temporal offsets in long recordings. On Table 7 it is also shown how much, in samples and in milliseconds, a given MUSE headset would drift away from the reference MUSE headset 1 after 3 hours of recording. Although for some headsets the drift is small for others like MUSE 4,5,6,8 and 10 the drift is in the order of hundreds of milliseconds. Although this might be small with respect to the total recording duration, it is a very significant drift when one aims at studying the alignment of brain signals. These drifts would have a very significant impact on studying whether the phase of brain signals are aligned at a given time instance. Given the frequency range of prominent brain activity, drifts in the order of 100s of milliseconds would lead to erroneous results. The same is true also for spectral power estimation, especially for higher frequencies with periods on the order of 10s of milliseconds. So, knowing the correct actual sampling frequencies is vital for studying inter-brain synchrony.

**TABLE 7.** Actual EEG Sampling Frequency of each MUSE headset. The corrected sampling frequency was calculated by aligning the Start and End Marker Pulses in the photodiode signal of each MUSE headset with those of MUSE headset 1, which served as the reference. In this way both the Start and End Marker Pulses were aligned across all MUSE headsets. The second data row shows how many samples(and msec in the parenthesis) is each MUSE headset drifted from the one expected from the assumed sampling frequency of 256 Hz, after a 3 hour recording. It is clear that the drift is from many headsets very substantial especially if one want to study phase - ocked phenomena or/and spectral power in the part of the typical brain spectrum.

|  | MUSE HEADSET ID |  |  |  |  |  |  |  |  |  |
| --- | --- | --- | --- | --- | --- | --- | --- | --- | --- | --- |
|  | 1 | 2 | 3 | 4 | 5 | 6 | 7 | 8 | 9 | 10 |
| Actual EEG Sampling Frequency Fsample (Hz) | 256.0000 | 256.0005 | 256.0007 | 256.0039 | 255.9895 | 255.9968 | 256.0009 | 256.0028 | 255.9977 | 255.9915 |
| Drift from assumed Fsample in a 3 hour recording. In samples (and msec) | na | 5.4<br>(21.1 msec) | 7.56<br>(29.5 msec) | 42.12<br>(164.5 msec) | -113.4<br>( -443 msec) | -34.56<br>( -135 msec) | 9.72<br>(37.9 msec) | 30.24<br>(118.1 msec) | -24.84<br>( -97 msec) | -91.8<br>( -358.6 msec) |

In summary the current experiment showed that the proposed RPi-based system for recording a large number of MUSE headsets is stable and consistent with extremely low loss of data in long recordings. Additionally, under these naturalistic conditions in cinema, the photodiode provided a marker signal that can be used to align the different MUSE headsets. Based on these markers the actual sampling frequency of each MUSE headset was identified.

### TESTING THE DELAYS OVER THE WI-FI NETWORK

This additional test recorded every 0.5 seconds the time difference between a message transmitted by the monitoring/control laptop and received by a RPi over a dedicated WI-FI.

This time difference is depicted for the entire recording of each of the 4 movies in Fig. 6. A thorough look on these plots shows that at times there are very large delays between message transmission and reception, in the order of even hundreds of seconds during some intervals. Even in cases where no such large delays occurred, for example during Movie 2 recording (Fig. 6B), the delays in many cases are in the order of 100s of milliseconds with a very high level of volatility. This volatility and the occurrence of very large delays at times show that alignment of MUSE data over a Wi-Fi network, such as in the case of using Lab Streaming Layer library for example, can be very problematic. That is why the demonstration that a photodiode can be used to provide unambiguous and stable alignment markers is an important contribution for the development of large-scale EEG recordings in audiences.

**Figure 6.**
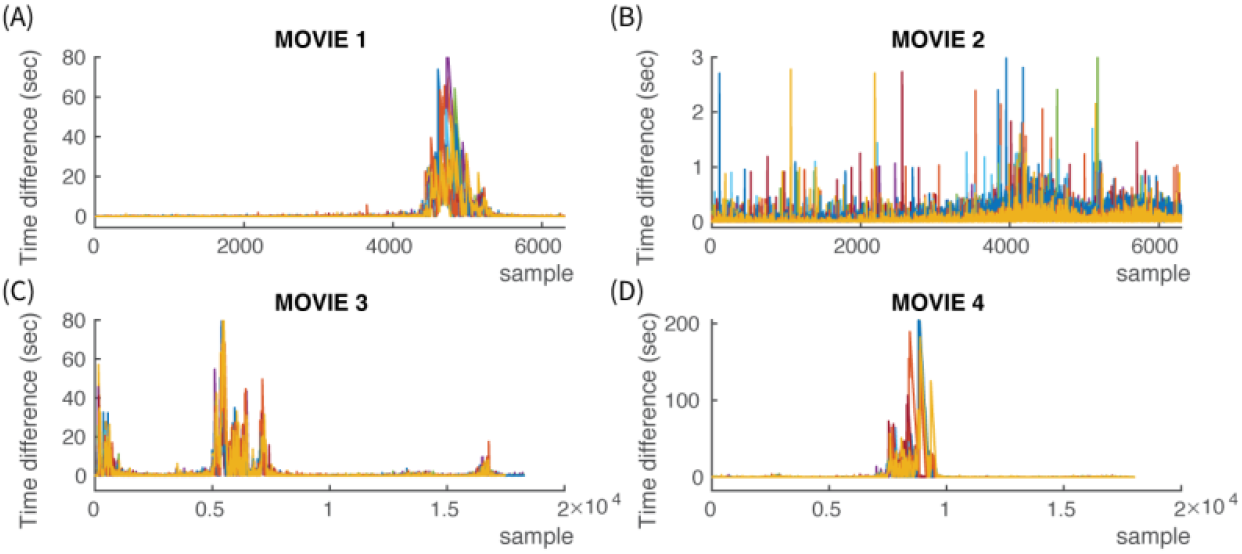
Delays Over Wi-Fi. Every 0.5 seconds the monitoring/control laptop was broadcasting its timestamp in the dedicated Wi-Fi network. This timestamp was recorded by each Raspberry Pi along with its own timestamp of the time that the broadcasted message was received. The main aim was to investigate in the 4 long recordings for the movies the range of delays that can occur in communication between devices through the Wi-Fi network. In (A) through (D) is depicted the time difference between the timestamp of message transmission and the timestamp of message reception over Wi-Fi during the recording in each of the 4 movies. During Movie 1 it can be seen in (A) that during a period towards the end of the recording there were very large delays in the order of 10s of seconds, probably representing router network instability. During Movie 2, as seen in (B), the delays remained bounded in much smaller values but still there was still high variability, with significant number of occurrences of delays near or larger than 1 second. During Movie 3 and Movie 4 it is seen, in (C) and (D) respectively, that there are intervals of very large delays of 10s or even 100s of seconds. Again these delays are probably caused by router connectivity issues during these periods.

## CONCLUSIONS

The current study makes important contributions towards the development of a scalable and low-cost off-the-shelve Hyper-scanning EEG system for recording audiences or large-groups. The system was tested successfully in realistic conditions with 10 MUSE headsets. Its scalability to larger numbers is supported by various factors. The first factor is that each MUSE headset is recorded in a dedicated Raspberry Pi. The second factor is that the developed software library, RPIMUSE, was developed based on a Bluetooth library, BLUEPY, specifically designed for RPi. This provided very stable connection between RPi and MUSE for very long periods with an extremely low percentage of data loss (which was not caused by loss of the Bluetooth connection). The third factor is the successful use of a photodiode for recording light patterns that can be used as alignment markers between the recorded data from different MUSE headsets. The availability of such unambiguous markers, independent of any network conditions, is vital for confidently aligning recorded data from a large number of EEG headsets. The combination of low-cost off-the-shelve hardware (MUSE, RPi, Photodiode) with stable, resilient software (BLUEPY, RPIMUSE) provides a unique system that can be scaled to large numbers for recording entire audiences.

Technically, the outlook for the next development steps is very promising. The current study was performed with the original MUSE-2016 headsets and with the Raspberry Pi 3+. Recently more advanced versions of these devices have been released. MUSE-2 is an upgraded version of the MUSE headset used here, with one more additional auxiliary port and with additional Photoplethysmography(PPG) sensors providing Heart rate and Breathing measurements with a sampling frequency of 64 Hz (InteraXon, 2022). The data communication through Bluetooth is the same as for the MUSE headsets used in this study, with more Bluetooth services available for subscription. So, the RPIMUSE library can be easily extended to record these new types of data. The Raspberry Pi Foundation has released the next generation, Raspberry Pi 4. This new version has much faster CPU, larger memory and better network adaptors. These characteristics are very likely to lead to data recording becoming even more stable, as the risk of processing bottlenecks causing data drops is reduced. Regarding triggering, the fact that the photodiode worked as a source of potential marker signals means that also other type of sensors or devices could be plugged in the auxiliary channel through the micro-USB port. One such could be a small battery-powered Radio Frequency receiver circuit, through which wireless triggers could be sent simultaneously to all headsets. In overall there are a lot of immediate technical improvements that can be made in the system tested in this study.

Regarding scalability, the recording of each MUSE on a dedicated Raspberry Pi is offering the opportunity of also developing scalable software. The open-source linux-based operating system offers great opportunities for developing software that can not only monitor a very large number of RPis but also control them. Such tasks are highly scalable, as the software technology already exists. One such example is the ZEROMQ (ZEROMQ, 2022) messaging library used in this study to estimate a simple network delay. This and other similar message exchange libraries can perform very sophisticated tasks in the coordination through messaging of a very large number of computers such as the RPi.

Although the MUSE headset has only 4 EEG sensors it can measure brain activity corresponding to a multitude of brain functions. In the current study it was demonstrated that the posterior MUSE electrodes can capture brain activity from early visual processing. Other studies have also demonstrated that it can measure ERP components of later visual processing (Krigolson et al., 2021; Krigolson et al., 2017). Multiple studies have demonstrated that it can be used to measure various aspects of brain function related to Stress (Asif et al., 2019; Phutela et al., 2022), Attention (Vortmann et al., 2022), Mindfulness (Hawley et al., 2021; Hunkin et al., 2021), Stroke (Wilkinson et al., 2020), Drowsiness (LaRocco et al., 2020), Emotion Classification (Raheel et al., 2019) and Fatigue(Krigolson et al., 2021; Ruyi et al., 2017). All these findings show that MUSE can measure to some extend both bottom-up and top-down processes of the brain.

The scalability of the system offers unprecedented opportunities for studying inter-brain synchrony in large groups. This is an area of Neuroscience research where only very few studies have been performed (Chen et al., 2022; Dikker et al., 2017) and where it is largely unexplored what are the brain processes that underlie collective and individual behavior in various types of large groups or crowds. Although many aspects of such behavior can be reduced and studied in very small groups or pairs, there are certain conditions that this cannot occur. One such case is the experience of an audience, especially in specific types of performances. Two of the most characteristic such cases, where the audience experience cannot be reduced to a small sample is the experience of an audience in sports and in theater. In this case the audience in its entirety is a vital part of the experience and the performance is planned with it as one of the most defining factors. A theatrical play or a football game performed without an audience is considered an oddity. The neural underpinnings that define the formation and role of an audience in these types of performances are largely unknown and this is an area of neuroscience research where systems such as the one presented in this study can provide an entry point.

